# Defining the Caprin-1 interactome in unstressed and stressed conditions

**DOI:** 10.1101/2021.01.06.425453

**Authors:** Lucas Vu, Asmita Ghosh, Chelsea Tran, Walters Aji Tebung, Hadjara Sidibé, Krystine Garcia-Mansfield, Victoria David-Dirgo, Ritin Sharma, Patrick Pirrotte, Robert Bowser, Christine Vande Velde

## Abstract

Cytoplasmic stress granules (SGs) are dynamic non-membranous foci containing translationally arrested mRNA and RNA binding proteins that form in response to a variety of cellular stressors. SGs may evolve into the cytoplasmic inclusions observed in many neurodegenerative diseases. Recent studies have examined the SG proteome by interrogating the interactome of G3BP1, a core SG protein. To gain further insight into the SG proteome, we employed an immunoprecipitation coupled with mass spectrometry approach of endogenous Caprin-1 in HeLa cells under unstressed or stressed conditions. Overall, we identified ~1,500 proteins that interact with Caprin-1. Interactors under stressed conditions were primarily annotated to the ribosome, spliceosome, and RNA transport pathways. We validated four Caprin-1 interactors that localized to arsenite-induced SGs: ANKHD1, Talin-1, GEMIN5, and SNRNP200. We also validated these stress-induced interactions in SH-SY5Y cells and determined that SNRNP200 also associated with osmotic and thermal induced SGs. Finally, we identified SNRNP200 in cytoplasmic aggregates in ALS spinal cord and motor cortex. Collectively, our findings provide the first description of the Caprin-1 protein interactome, identify novel cytoplasmic SG components, and reveal a SG protein in cytoplasmic aggregates in ALS patients. Proteomic data collected in this study are available via ProteomeXchange with identifier PXD023271.

## Introduction

In response to environmental stress, eukaryotic cells form non-membranous condensates, termed stress granules (SGs). SGs form upon environmental insults that inhibit global translation such as thermal, oxidative and osmotic stress, and contain untranslated mRNAs as well as a multitude of RNA binding proteins (RBPs) (1, 2). SG formation is a transient cellular response and SG dissociation upon stress dissipation is equally important as SG assembly. Upon removal of stress, the release of stalled transcripts, ribosomal components, and other translation initiation machinery that are sequestered in SGs, is vital for cell survival (3).

Several RBPs that are associated with SG formation bear low-complexity domains that facilitate liquid-liquid phase separation (LLPS) and condensate formation (4). Upon prolonged stress, SGs can transform into more solid aggregates which are irreversible (5). Multiple neurodegenerative diseases are characterized by the presence of pathological aggregates (6) and defects in SG dynamics are thought to seed such pathological aggregates (7). A variety of RBPs associated with SGs contain disease-causing mutations that are linked to amyotrophic lateral sclerosis (ALS), Alzheimer disease (AD), and frontotemporal dementia (FTD) (8–12).

There has been considerable effort invested to identify and understand the proteomic (13, 14) and transcriptomic (15, 16) composition of SGs (14, 17–20). Most of these studies have used G3BP1, a core SG protein, to define the SG proteome (19). It is plausible that some proteins localize to SGs independent of G3BP1 and thus, would not be detected as a SG component by these prior studies. Furthermore, many of these prior studies used over-expression of SG proteins to identify protein interactors, and this may perturb normal protein stoichiometry within SGs, potentially leading to erroneous interactions and/or mis-recruitment of proteins to RNP granules. Unravelling the SG proteome is essential to further understanding disease mechanisms involving SG dynamics.

In this study, we define the Caprin-1 protein interactome in unstressed and sodium arsenite (SA) induced oxidative stress conditions. We utilized immunoprecipitation coupled to mass spectrometry (IP-MS) to identify ~1,500 Caprin-1 protein-protein interactions (PPIs) in either unstressed conditions or following SA stress. Our results provide insight into the multitude of Caprin-1 mediated biological functions, including RNA transport and translation. Validation of select interactors resulted in annotating novel SG localized proteins in HeLa cells. Among them, SNRNP200 was found in SGs induced by oxidative, thermal, as well as osmotic stress in both non-neuronal and neuronal-like cells. Furthermore, we found SNRNP200 in cytoplasmic aggregates within motor neurons of both spinal cord and motor cortex in ALS patients. Our work identifies Caprin-1 PPIs and characterizes one that is also a component of SGs assembled in neuronal-like cells.

## Experimental section

### Cell culture and stress

HeLa cells were maintained in complete media containing DMEM (ThermoFisher) and 10% fetal bovine serum (FBS) (Gibco, Life Technologies) and 1% Penicillin-Streptomycin (ThermoFisher) at 37°C. SH-SY5Y cells were grown in DMEM/F12 media (ThermoFisher) containing 10% FBS (Gibco, Life Technologies). SH-SY5Y cells were differentiated using sequential treatments of retinoic acid (RA) (Cedarlane) and brain-derived neurotrophic factor (BDNF) (Sigma) (21). Briefly, cells seeded on collagen coated plates were maintained in 10 μM RA containing media for 5 days with daily media changes. Thereafter, cells were grown in 50 ng/mL BDNF containing media for 12 days. Stress was performed on day 12 and cells were fixed immediately with 4% paraformaldehyde (PFA, FD NeuroTechnologies) for staining.

Cells were seeded, maintained, and stressed on glass coverslips. To induce oxidative stress, cells were treated with 0.5 mM of SA (Sigma) for 30 mins at 37°C. The media was replaced with fresh media and incubated at 37°C for 60 min to initiate recovery after SA stress (22, 23). Heat shock was performed by incubating cells at 43°C for 30 mins. Osmotic stress was performed by incubating the cells in 0.4 M sorbitol (Sigma) containing media at 37°C for 2 hrs. Following stress, all cells were immediately fixed with 4% PFA.

### Immunofluorescent staining

Fixed cells were permeabilized with 0.1% triton X-100 solution in 1X PBS. Cells were then blocked using Super Block (Scytek) and subsequently incubated overnight at 4°C with primary antibodies diluted in Super Block. Cells were washed and incubated with Alexa-Fluor secondary antibodies for 1 hour at room temperature. All antibodies used in this study are shown in Table S1. Nuclei were labelled with either 41,6-diamidino-2-phenylindole (DAPI, ThermoFisher) or Hoechst 333432 (ThermoFisher). Cover slips were mounted on slides using ProLong antifade reagent (Invitrogen). Images were acquired using a 63X oil lens on Zeiss LSM 710 (Zeiss) or a Leica SP5 (Leica Microsystems) confocal microscope. Line scan analyses were performed to assess co-localization using Image J V1.52a (24).

### Immunostaining of human post-mortem ALS tissues

Paraffin-embedded spinal cord and motor cortex tissue sections from 5 ALS, 2 disease controls (DCs) and 2 non-neurologic diseased controls (NNDCs) were obtained from the Barrow Neurological Institute and Target ALS post-mortem tissue bank cores (subject demographics listed in Table 1). Participants of the post-mortem tissue bank cores provided IRB approved informed consent for the collection of post-mortem tissues. Immunohistochemistry (IHC) was performed as previously described (25). Tissue sections were de-paraffinized and rehydrated, and antigen retrieval was performed using the Antigen Retrieval Citra Solution (pH 6) (BioGenex). Tissues were blocked using Super Block supplemented with Avidin (Vector Labs) and incubated with rabbit anti-SNRNP200 antibody (Sigma) diluted in Super Block supplemented with biotin (Vector Labs) overnight. An anti-rabbit biotinylated IgG secondary antibody (Vector Labs) was subsequently added, and immunostaining was visualized using the Vectastain Elite ABC reagent (Vector Labs) and Vector ImmPACT NovaRED peroxidase substrate kit (Vector Labs). Slides were counterstained with hematoxylin (Sigma). Images were acquired using an Olympus BX40 microscope and quantified using Image J (25). Details on antibodies used for IHC are listed in Table S1.

**Table 1:**
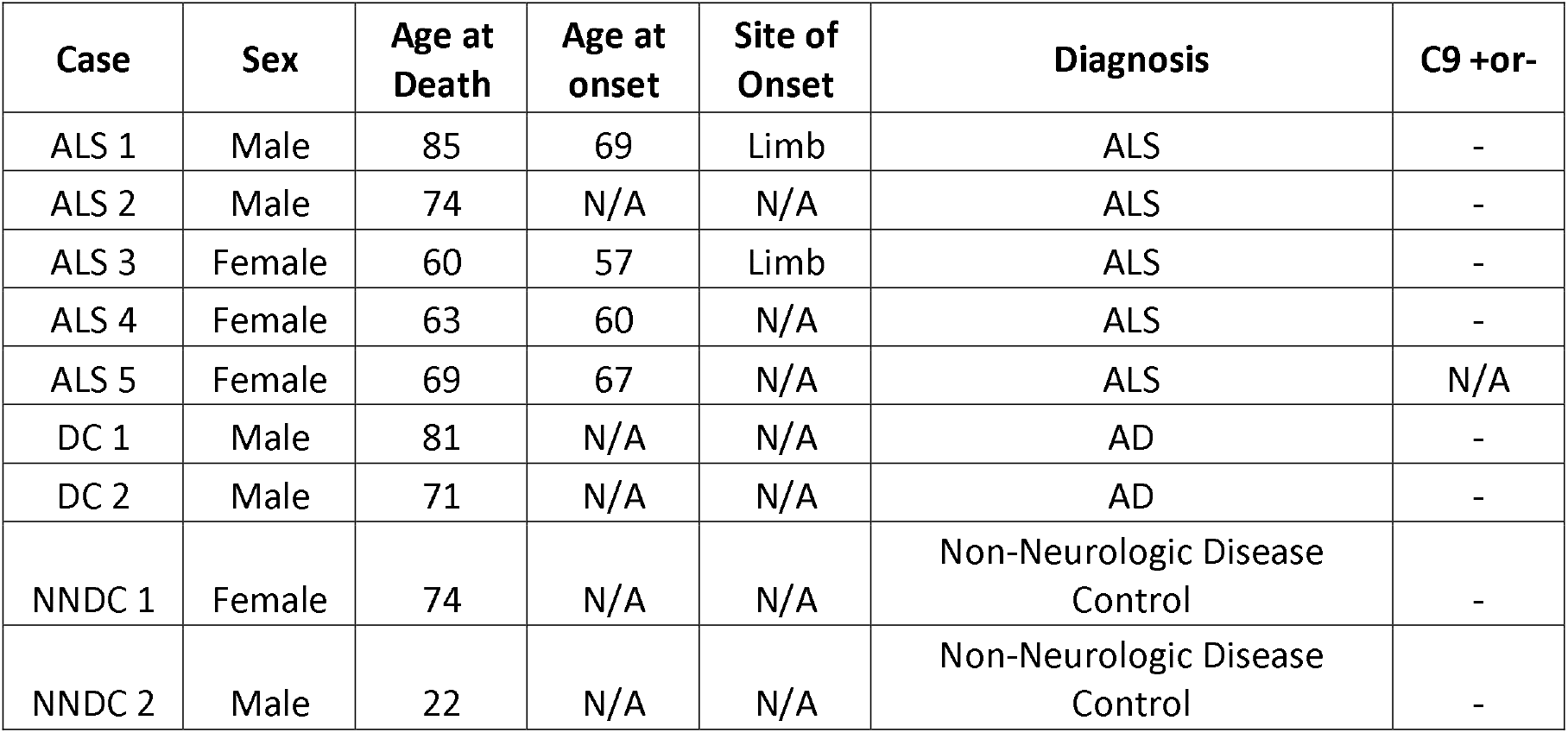
Demographics of patients used for SNRNP200 immunohistochemistry. N/A – Not assessed

### Cell lysis and Immunoprecipitation for Caprin-1

HeLa cells were grown in 100 mm dishes until 95% - 100% confluent. Following stress, cells were crosslinked with 0.1 % formaldehyde and quenched with 1.25 M glycine. Cells were subsequently scraped into IP lysis buffer (50 mM HEPES pH 7.2, 150 mM NaCl, 0.1 % NP-40, 10% Glycerol, 2 mM EDTA, 2 mM EGTA, 1 mM DTT, Protease inhibitor (Sigma Aldrich) and Ribolock (ThermoFisher), and incubated on a rotator at 4°C for 20 mins. Lysates were centrifuged at 10,000 x g for 15 mins at 4°C and the supernatant was collected. Protein content of the supernatants were determined using the Bradford protein assay (BioRad). Each immunoprecipitation was carried out in quadruplicate. 1 μg of rabbit anti-Caprin-1 (Proteintech) or rabbit anti-IgG (Jackson Immuno Research Laboratories) was added to magnetic protein A Dynabeads (ThermoFisher). Immunoprecipitation was performed with an input of 400 μg of total protein for 1 hr at room temperature. After IP, the beads were washed 6 times with IP lysis buffer (without DTT, protease inhibitors, and Ribolock) and stored at −20°C.

### Protein digestion and peptide sample preparation

The immunoprecipitated protein was eluted from the beads with 1X Laemmli buffer (BioRad) by heating at 95°C for 10 mins. Samples were resolved via gel electrophoresis and stained with Bio safe Coomassie stain (BioRad) for 1 hr and destained in double deionized water overnight. Each lane was excised into three individual fractions with IgG heavy (50 kDa) and light (25 kDa) chains excluded. The fractions were cut into 1-2 mm^3^ cubes and further de-stained, washed and dried. Proteins were reduced by incubating with 10 mM DTT (ThermoFisher) for 30 mins at 60°C and alkylated with 55 mM iodoacetamide (ThermoFisher) for 30 mins at room temperature using a series of hydration and dehydration steps. Digestion was performed using Trypsin gold (Promega) at 1:20 enzyme to protein ratio and incubated overnight at 37°C. After digestion, trypsin was inactivated by adding trifluoroacetic acid (ThermoFisher) to a final concentration of 0.5% (v/v). Peptides were extracted from the gel pieces by hydration and dehydration and further clean-up was performed using C18 StageTips as previously described (26, 27).

### LC-MS/MS analysis

Data acquisition was performed on a Thermo Orbitrap Fusion Lumos interfaced with a U3000 RSLCnano UHPLC operating at a flow rate of 300 nL/min, with Solvent A (water, 0.1% FA) and Solvent B (acetonitrile, 0.1% FA). Peptides in 5 μL of loading solvent (98% water, 2% acetonitrile, 0.1% FA) spiked with 25 femtomoles of synthetic peptides (Pierce Retention Time Calibrator Mix, ThermoFisher) were directly loaded on a 15 cm C18 EasySpray column (ES800, ThermoFisher) maintained at 45°C. Peptides were separated over 60 mins using the following gradient: 2-19% B in 42 mins, 19-45% B in 6 mins and then to 90% B in 0.5 min, isocratic at 90% B for 1 min followed by return to initial conditions in 0.5 min and column equilibration for 10 mins. Data-Dependent Acquisition (DDA) of eluted peptides was performed using top-speed mode with a cycle time of 3 secs. MS1 scans were acquired in the Orbitrap at a resolution of 120K, with a mass range of 400-1500 m/z, keeping the AGC target at 4E5 and with a max ion injection time to 50 millisecs. Most abundant precursor ions with a charge state between 2 and 7 were selected with an isolation window of 1.6 Da and fragmented using High Energy Collision Dissociation (HCD), followed by detection in the ion trap. A dynamic exclusion of 60 secs was used to prevent resampling of the same precursors during the elution of chromatographic peaks.

### Data processing

Mass spectra were aligned across replicate bands for each IP and condition, within Progenesis QI for Proteomics v4.1.6 (Nonlinear Dynamics, A Waters Company) using default parameters for automated processing. Aligned and filtered spectra were searched against a human database (Swissprot/UniprotKB, 2017) in Mascot v2.6 (Matrix Science) with trypsin cleavage rules, and a maximum of 2 missed cleavages. Allowed dynamic modifications were oxidation (methionine) and acetylation (N-term, lysine). Cysteine carbamidomethylation was set as a fixed modification. Matched spectra were imported into Progenesis QI and only peptides with a Mascot homology score >21 were retained. Gel bands were recombined for protein identification and quantitation.

Proteins were first filtered for their presence in >75% of samples and then normalized using the cBioconductor DEP package in R (28). Average probabilities of interaction (AvgP) were calculated using the SAINTexpress algorithm (29) against IgG controls. High confidence interactors (AvgP >0.7 and presence in 3 out of 4 replicates) were compared between the stressed and unstressed cell lines to determine unique Caprin-1 interactors for each condition. The mass spectrometry proteomics data have been deposited to the ProteomeXchange Consortium via the PRIDE partner repository (30) with the dataset identifier PXD023271. Reviewer account details: Username: Password: 5XWxNAlT

### StringDB, gene ontology, pathway analysis

High confidence proteins unique to either the stressed or unstressed condition were imported into StringDB to reveal protein clusters and associations previously annotated in literature. Networks were created from all interaction evidence in the String database, and filtered for a String score >0.7, with unconnected nodes removed from the figure. Gene ontology and KEGG pathway analysis was performed using DAVID V6.8 with default settings (31).

### Statistical analyses

Statistical analyses were performed using GraphPad Prism 8. For the motor neuron quantification, a Mann-Whitney test was performed to compare differences between ALS and controls with p < 0.05 considered significant.

## Results

### Caprin-1 immunoprecipitation and mass spectrometry

In order to interrogate the interactome of endogenous Caprin-1 (Figure 1A), we employed immunoprecipitation (IP) of endogenous Caprin-1 from HeLa cells exposed to SA stress and compared against unstressed controls. We followed a paradigm of 30 mins of acute SA stress followed by 60 mins of recovery which captures the secondary assembly of SGs, as previously described (23). After stress, cells were cross-linked with formaldehyde to preserve all interactions. Immunoblot using anti-Caprin-1 antibody confirmed enrichment of Caprin-1 in the Caprin-1 IP lanes but not in the control IgG IP lanes (Figure 1B). The IP fractions were separated by SDS page, and subjected to reduction, alkylation and trypsin digestion and subsequently analyzed by data-dependent LC-MS/MS. Overall, we identified ~1500 proteins for Caprin-1 IPs and ~500 proteins from the IgG control pull-downs (Figure 1C, 1D, Table S2) with known SG components also being identified in both the stressed and unstressed Caprin-1 IPs (e.g. Ataxin-2, USP10, and TIAR).

**Figure 1:**
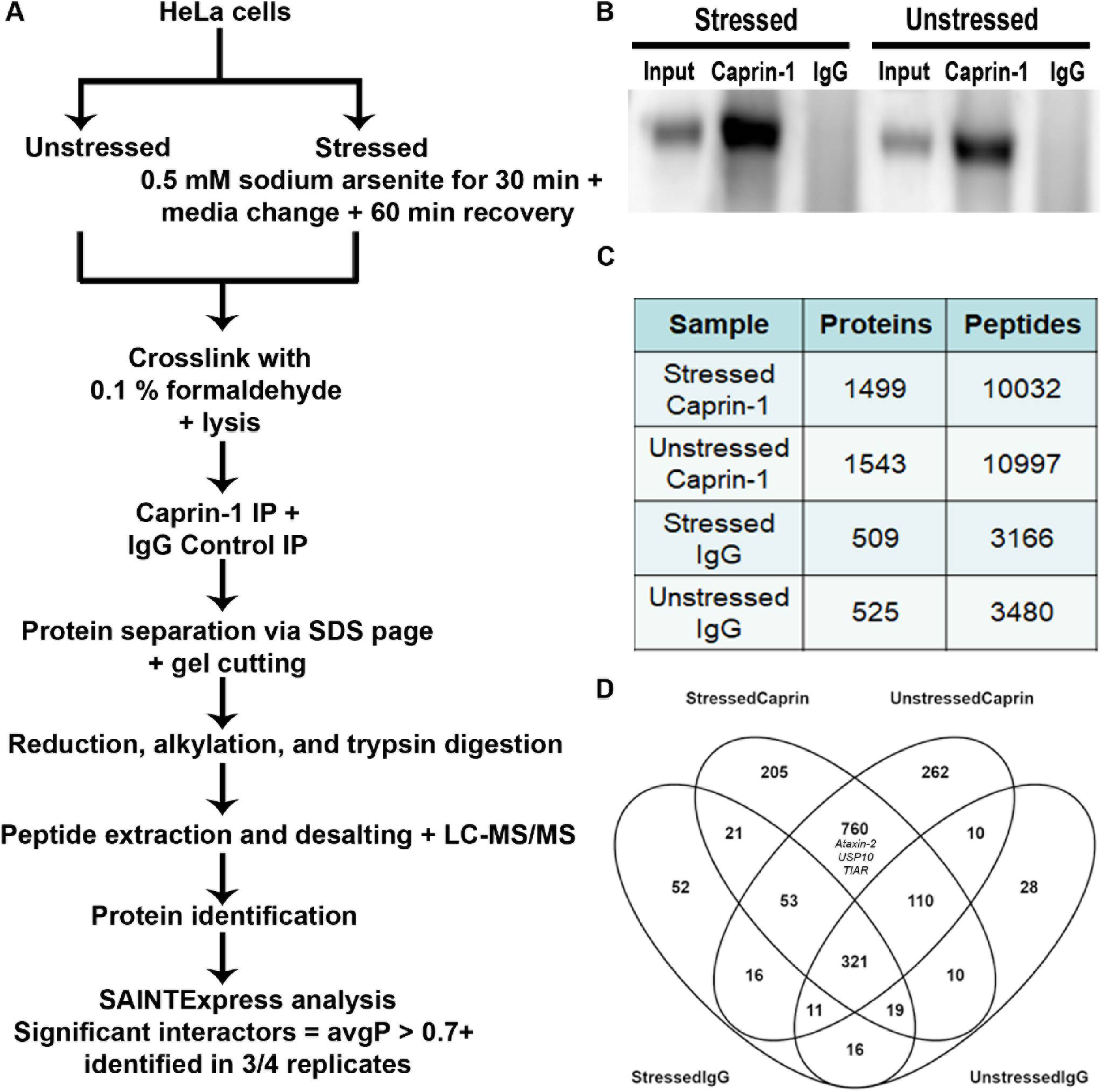
Caprin-1 IP-MS methods and analysis of the interactome. A) Schematic of the methodology employed in this study. B) Western blot of Caprin-1 and IgG IP fractions from lysates derived from both stressed and unstressed conditions demonstrating the presence of Caprin-1 only in the Caprin-1 IP and not in the IgG control. C) Summary of the proteins and peptides identified by LC-MS/MS from quadruplicate IP experiments per condition. D) Venn diagram demonstrating the overlap of all of the proteins identified across all comparisons.

To further characterize the interactome of Caprin-1 under stressed and unstressed conditions, we employed SAINTexpress, an algorithm premised on label free IP-MS experiments to detect PPIs (32) that we have successfully used to define the protein interactome for other disease related proteins (33, 34). Proteins identified in at least 3 of the 4 replicates and demonstrating an average probability of interaction (AvgP) greater than 0.7 were considered high confidence interactors of Caprin-1 (Figure 1A). Based on this criterion, we identified 281 and 326 proteins that interact with Caprin-1 under stressed and unstressed conditions respectively, with 38% (166/441) overlap between the two conditions (Figure 2A, Table S3). Of the 166 commonly identified proteins in both stressed and unstressed conditions, 10 proteins have been previously reported to play a functional role in stress granules. This includes core SG proteins like G3BP1 and G3BP2-2 (also known as G3BP2b) which reflects that pre-stress interactions between SG proteins exist and perhaps is the reason for rapid dynamics in SG formation at the onset of stress. Other SG components like USP10, FMR1, Ataxin2-like (ATXN2L) and UPF1 are also pre-stress interactors of Caprin-1. Multiple RNA helicases like DDX1, DDX3X, and DDX5 which have known roles in RNP granules (including SGs and P-bodies) are also pre-stress interactors of Caprin-1 (Figure 2A). CNOT7, which is localized to P-bodies and participates in mRNA decay, was also detected as a Caprin-1 interactor in both conditions.

**Figure 2:**
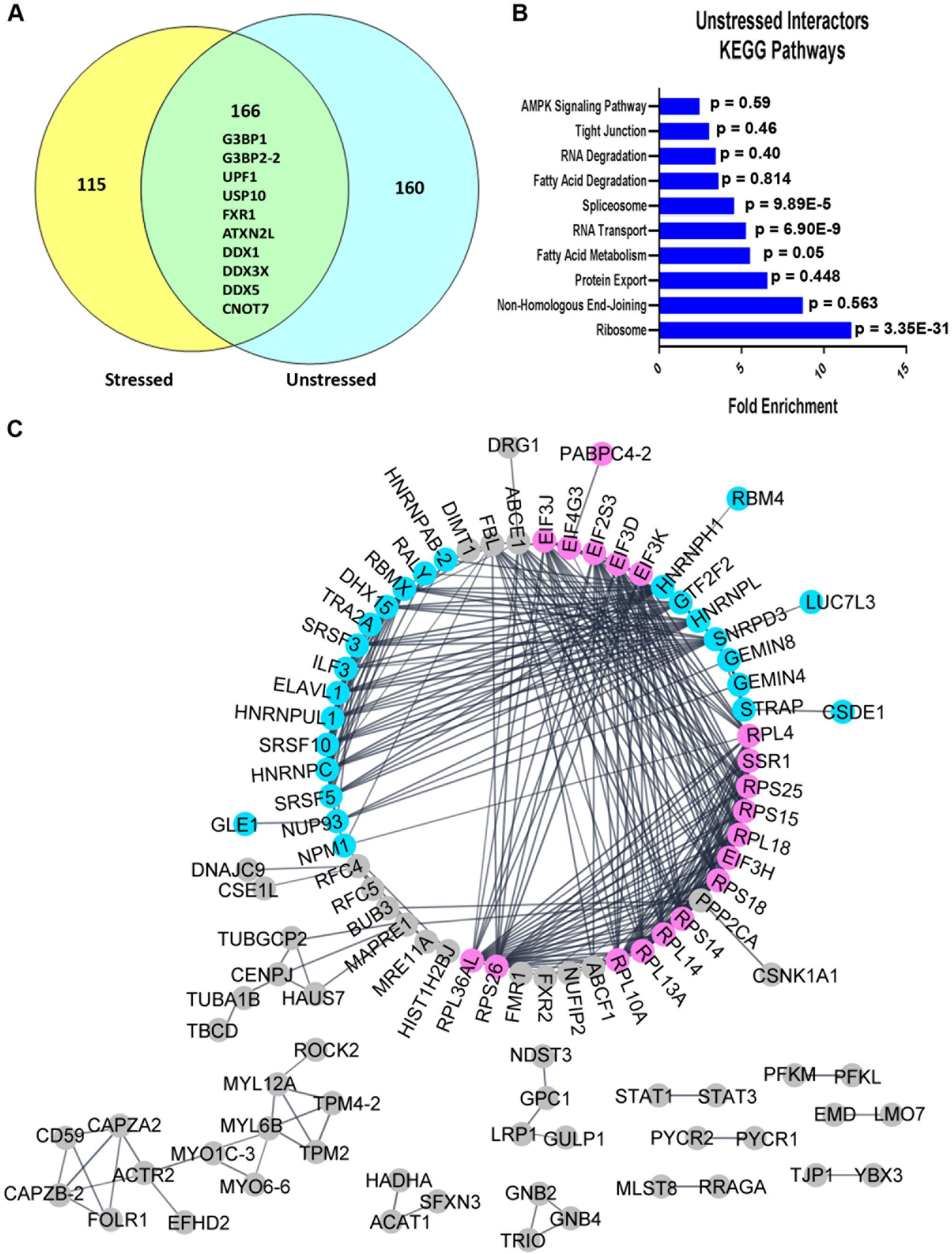
Analysis of the unstressed interactome of high confidence Caprin-1 interactors revealed associations with ribosomes, RNA transport, and spliceosome pathways. A) Venn Diagram comparing high confidence interactors of Caprin-1 in both stressed and unstressed conditions, as determined by SAINTexpress. A 38% overlap was observed between conditions. Notable proteins common between the two conditions are indicated. B) KEGG pathway analysis of the unstressed Caprin-1 interactors. C) StringDB analysis demonstrating protein-protein interactions between the unique interactors of Caprin-1 in unstressed conditions, with distinct groups annotated as RNA binding proteins (pink) and splicing factors (blue). All other interactors are labeled in gray.

### Caprin-1 interactome in unstressed conditions

To identify biological pathways in which Caprin-1 functions in unstressed conditions, we employed KEGG pathway analysis using all 326 high confidence interactors of Caprin-1 and found significant enrichment of pathways related to ribosomes, RNA transport, and the spliceosome suggesting that Caprin-1 participates in these pathways in unstressed conditions (Figure 2B, Table S4). Gene ontology enrichment analyses of the top 15 biological processes and molecular functions revealed similar findings as KEGG pathway analysis, with terms broadly related to ribosomes and splicing (Figure S1). More specifically, this analysis suggested a functional role of Caprin-1 in translation initiation, mRNA splicing, and RNA binding (Figure S1A, S1B). To investigate known PPIs between Caprin-1 and the unstressed interactors, StringDB analysis was employed on 160 uniquely interacting proteins in the unstressed conditions (Figure 2A). Within the largest interaction network, we identified two groups which correspond to splicing factors (highlighted in blue) and ribosome and translation related proteins (highlighted in pink) that interact with Caprin-1 (Figure 2C). This complementary analysis recapitulated the gene ontology analysis, re-affirming that the primary functions of Caprin-1 are associated with translation and RNA splicing.

### Caprin-1 interactome following sodium arsenite stress

We next interrogated the Caprin-1 interactome in SA stress conditions to understand how the interactome of Caprin-1 changes when it predominantly localizes to SGs (281 proteins in Figure 2A). KEGG pathway analysis demonstrated similar enrichment of pathways as compared to the unstressed interactors of Caprin-1 where the stress-dependent interactors were annotated to the ribosome, RNA transport, and spliceosome, and additionally to protein export (Figure 3A, Table S5). Interestingly, nearly half (131/281) of the high confidence interactors were annotated to known pathways (Figure 3B, Table S5). As Caprin-1 is primarily localized to SGs upon stress (35) and SGs are known to modulate translation (3), it was not surprising that several translation-related proteins and ribosomal subunits were among Caprin-1 interactors. StringDB analysis of the unique stress-dependent interactors also demonstrated two interaction networks corresponding to splicing factors and ribosomal proteins as Caprin-1 interactors (Figure S2).

**Figure 3:**
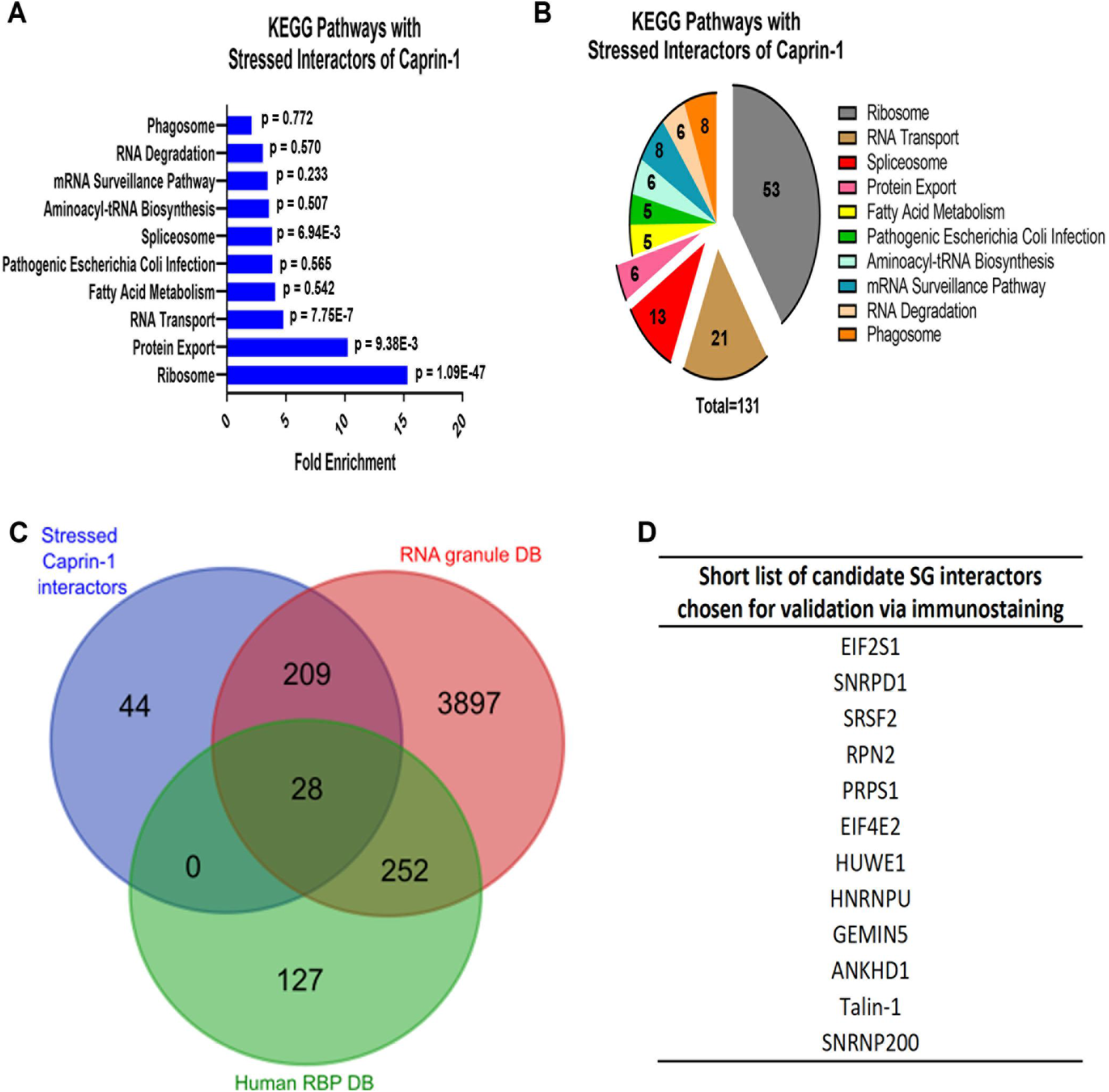
Analysis of the stressed Caprin-1 interactome and shortlisted hits for validation. A) Pathway analysis of Caprin-1 interactors in stressed conditions (326 proteins). B) Pie chart demonstrating the number of Caprin-1 stressed interactors annotated to each pathway. C) Venn Diagram demonstrating the overlap between the stress-dependent Caprin-1 interactors (blue) and RNA binding proteins from Human RBPs (green, http://rbpdb.ccbr.utoronto.ca/) and RNA Granule DB (red, http://rnagranuledb.lunenfeld.ca/). D) Short list of 12 candidate Caprin-1 interactors derived from pathway and RBP analyses that were further interrogated by immunostaining to assess co-localization with Caprin-1.

Prior studies have shown that Caprin-1 interacts with G3BP1 in SGs (14, 19, 35) suggesting that there may be some degree of overlap among their respective interactomes. Therefore, we compared our stress dependent Caprin-1 interactors (281 proteins) with previously published SG proteomes (Figure S3). BioID based profiling of Caprin-1 interactors demonstrated only 14 interactors that overlapped with our Caprin-1 interactors. Interestingly, of these 14, USP10, FXR1, UPF1 are known SG components (Figure S3A). There was also a minor overlap with the BioID G3BP1-interactome and notable SG components such as PABPC1, PABPC4, ATXN2L were common interactors of both Caprin-1 and G3BP1 (Figure S3B). While there were only 14 proteins that overlapped between our Caprin-1 interactome and either BioID-Caprin-1 or BioID-G3BP1 interactomes, four proteins were common to all the three datasets (PABPC4, FXR1, UPF1 and USP10) (Table S6). Comparison of our Caprin-1 interactors with recent publications probing the SG proteome with GFP-G3BP1 also showed minimal overlap, including multiple RBPs such as SYNCRIP, IGF2BP1, and HNRNPA3 (Figure S3C, Table S6). Lastly, only 28 proteins overlapped between our Caprin-1 dataset and that obtained using Apex-mediated proximity labelling of G3BP1 as bait, with notable translation-related SG components such as EIF3L, EIF3A, TAF15 being common (Figure S3D, Table S6).

As Caprin-1 is an integral SG protein that may interact with multiple RBPs, we also compared our list of high confidence interactors with known RBPs listed in two independent databases: Human RBP DB v1.3.1 (http://rbpdb.ccbr.utoronto.ca/) and RNA Granule DB v1 (http://rnagranuledb.lunenfeld.ca/) (Figure 3C, Table S7). Our results demonstrate 84% overlap (237/281) between proteins within our stressed candidate list (281 proteins from Figure 2A) and known RBPs. Combining protein lists from both the significantly enriched KEGG pathways as well as those that overlap between our interactors in stressed conditions and known RBPs, we shortlisted 12 Caprin-1 interactors that had not been previously associated with SGs to assess their localization in SGs (Figure 3D). We chose to focus on RBPs with no prior evidence of localization to SGs as well as those that annotated to the spliceosome and RNA transport pathways since RNG105 (*Xenopus* ortholog of human Caprin-1) has been previously shown to concentrate in ribosomal-associated granules (36) and the presence of various translation-related proteins to SGs is widely reported (37).

### Validation of Caprin-1 interactors in HeLa cells

To assess if our candidate Caprin-1 interacting proteins co-localize with Caprin-1 labelled SGs, we performed double label immunofluorescence microscopy in HeLa cells following SA stress. Among the 12 candidates we prioritized for validation, ANKHD1, Talin-1, GEMIN5, and SNRNP200 demonstrated co-localization with Caprin-1 labeled SGs following exposure to SA (Figure 4). While one prior study has identified ANKHD1 as an interactor of Caprin-1 (18), this is the first report of ANKHD1 being visualized in SGs by immunostaining. The remaining 8 candidates failed to co-localize with SGs (Figure S4). As SG composition can vary with different stressors (38), we assessed the potential of ANKHD1, Talin-1, GEMIN5, and SNRNP200 to co-localize with SGs induced by thermal (heat shock) and osmotic (sorbitol) stressors. Under these conditions, both ANKHD1 and SNRNP200 (Figure 5A, 5B) co-localized with Caprin-1 SGs while Talin-1 and GEMIN5 did not (Figure 5C, 5D).

**Figure 4:**
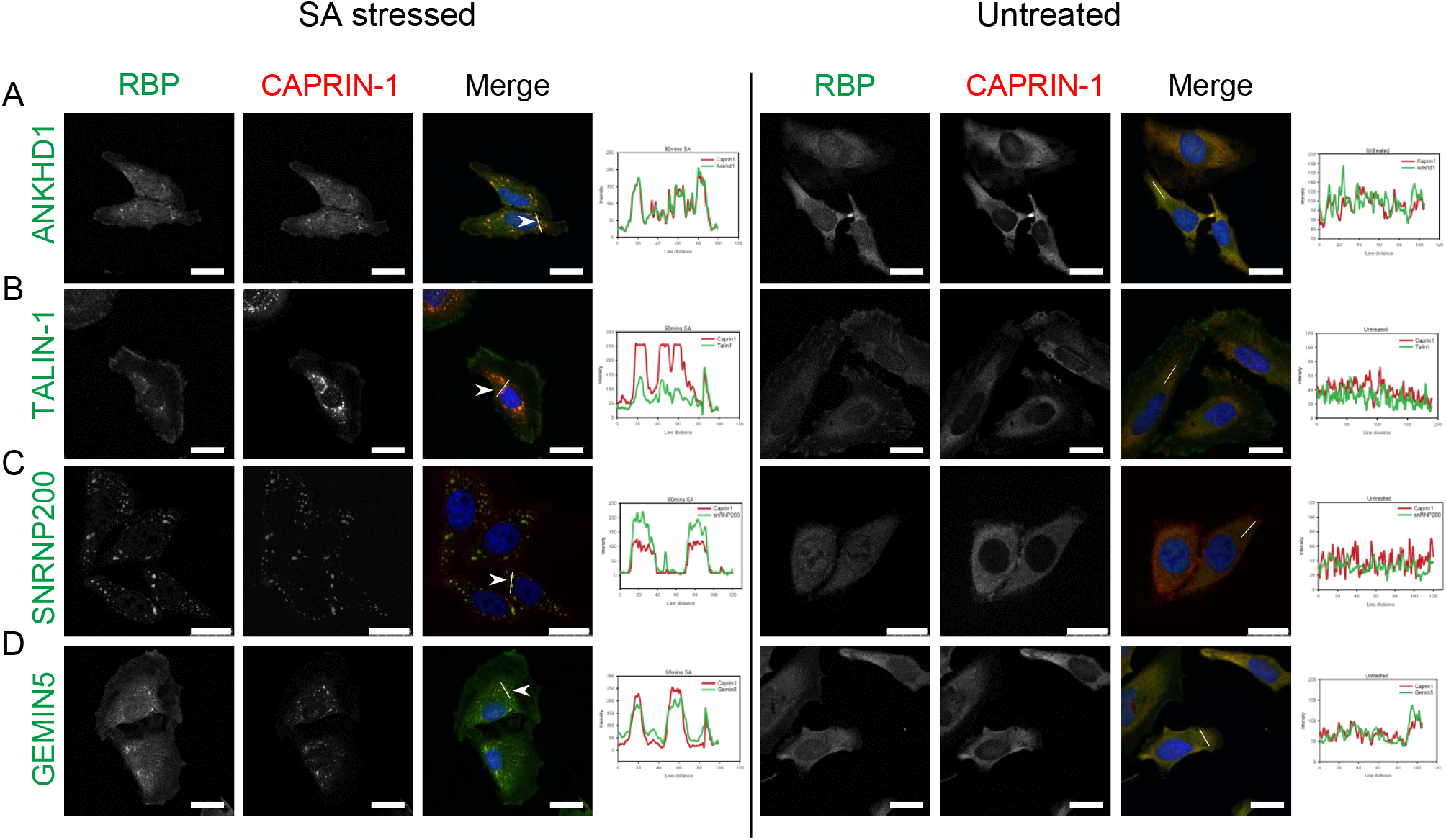
Arsenite stress induced Caprin-1 interactors; ANKHD1, Talin-1, SNRNP200, and GEMIN5 co-localize with SGs in HeLa cells. HeLa cells were stressed with either 0.5mM SA (left panel) or, left untreated (right panel). Representative images of immunofluorescent labelling for Caprin-1 and A) ANKHD1, B) Talin-1, C) SNRNP200, and D) GEMIN5. Nuclei were visualized with Dapi. Scale bar, 25 μm. Line scan-based co-localization analysis is plotted as histograms.

**Figure 5:**
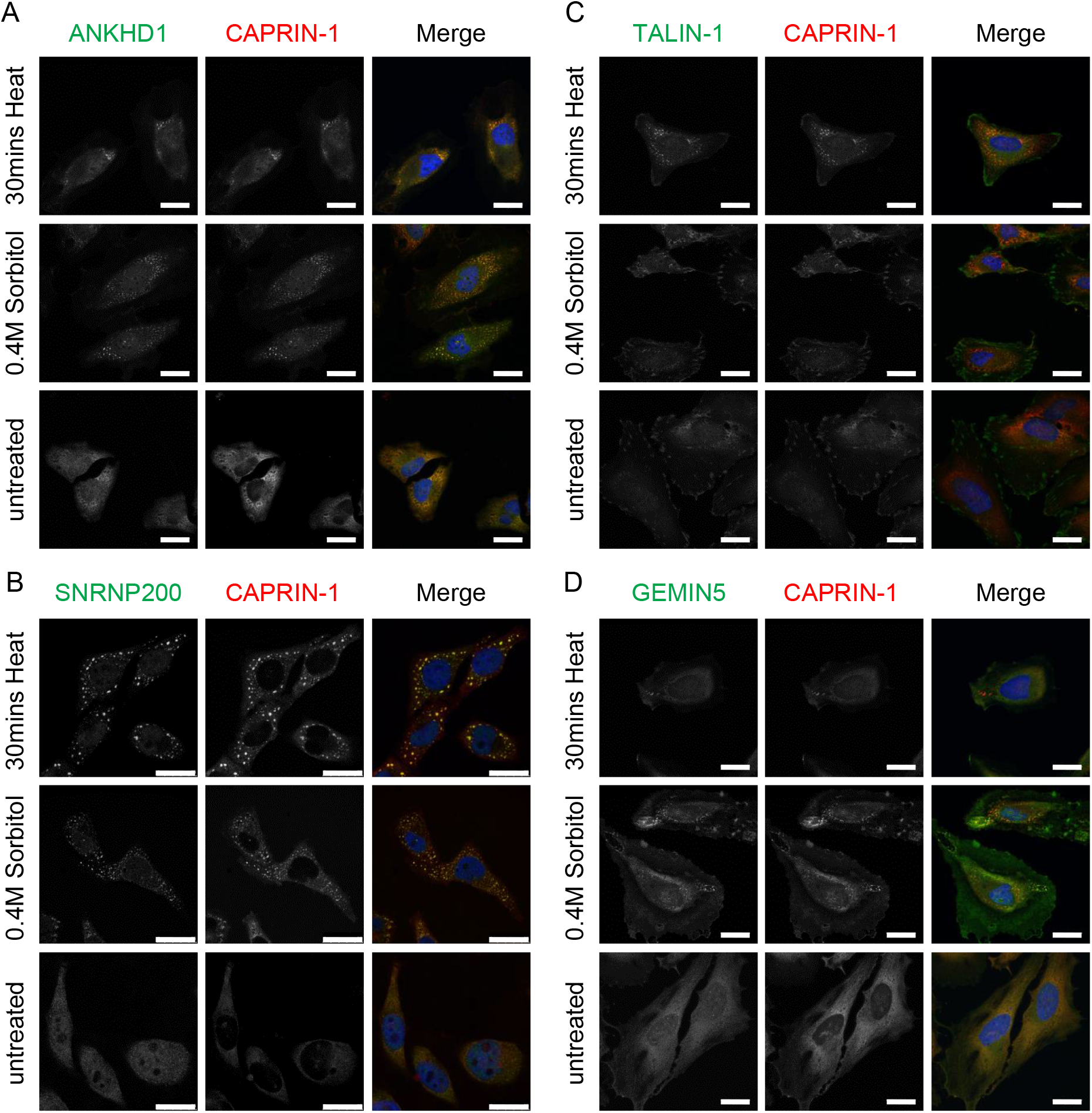
Caprin-1 interactors; ANKHD1 and SNRNP200 co-localize with SGs in HeLa cells subjected to thermal and osmotic stressors. HeLa cells were stressed with either 30 mins heat stress at 43°C or 0.4M sorbitol for 2 hrs or, left untreated. Representative images shown after co-labelling for Caprin-1 and A) ANKHD1, B) SNRNP200, C) Talin-1, and D) GEMIN5. Nuclei were visualized with Dapi. Scale bar, 25 μm.

### SNRNP200 co-localizes with Caprin-1 in SGs in neuronal-like cells

It is reported that SG components can differ among various cell types, with neuronal cells having a more diverse SG composition (19). To determine the propensity of the four novel Caprin-1 interactors (ANKHD1, Talin-1, GEMIN5, and SNRNP200) to co-localize with SGs in a neuronal-like context, we utilized the SH-SY5Y human neuroblastoma cell line. SNRNP200 co-localized with Caprin-1 labelled SGs following SA or heat shock induced stress (Figure 6A). In contrast, ANKHD1, Talin-1 and GEMIN5 did not form any punctate structures in SH-SY5Y cells during thermal or SA stress (Figure S5A-C). This further demonstrates the variability of SG composition between cell types. As the SG composition may vary in terminally differentiated cells, we also assessed SNRNP200 in SH-SY5Y cells differentiated with BDNF. We validated the differentiation of SH-SY5Y cells using the neuronal marker Tubb3/Tuj1 (Figure S5D). Indeed, we found SNRNP200 co-localizes with Caprin-1 labelled SGs in terminally differentiated SH-SY5Y cells (Figure 6B). Line scans to evaluate co-localization indicate clear overlap of both the Caprin-1 and SNRNP200 signals following acute thermal or SA stress (Figure 6B, right panel; Table 2).

**Table 2:** Summary of immunostaining to assess co-localization of the candidate RBPs with SGs.

| Candidate RNA binding protein | In HeLa cells |  |  | In SH-SY5Y cells |  | Differentiated SH-SY5Y |  |
| --- | --- | --- | --- | --- | --- | --- | --- |
|  | Sodium arsenite stress | 43°C heat stress | 0.4M Sorbitol stress | Sodium arsenite stress | 43°C heat stress | Sodium arsenite stress | 43°C heat stress |
| HUWE1 | No | N/A | N/A | N/A | N/A | N/A | N/A |
| HNRNPU | No | N/A | N/A | N/A | N/A | N/A | N/A |
| ANKHD1 | Yes | Yes | Yes | No | No | No | No |
| Talin-1 | Yes | No | No | No | No | No | No |
| GEMIN5 | Yes | No | No | No | No | No | No |
| SNRPD1 | No | N/A | N/A | N/A | N/A | N/A | N/A |
| EIF2S1 | No | No | N/A | N/A | N/A | N/A | N/A |
| PRPS1 | No | N/A | N/A | N/A | N/A | N/A | N/A |
| RPN2 | No | N/A | N/A | N/A | N/A | N/A | N/A |
| SRSF2 | No | N/A | N/A | N/A | N/A | N/A | N/A |
| SNRNP200 | **Yes** | **Yes** | **Yes** | **Yes** | **Yes** | **Yes** | **Yes** |

N/A: not assessed

**Figure 6:**
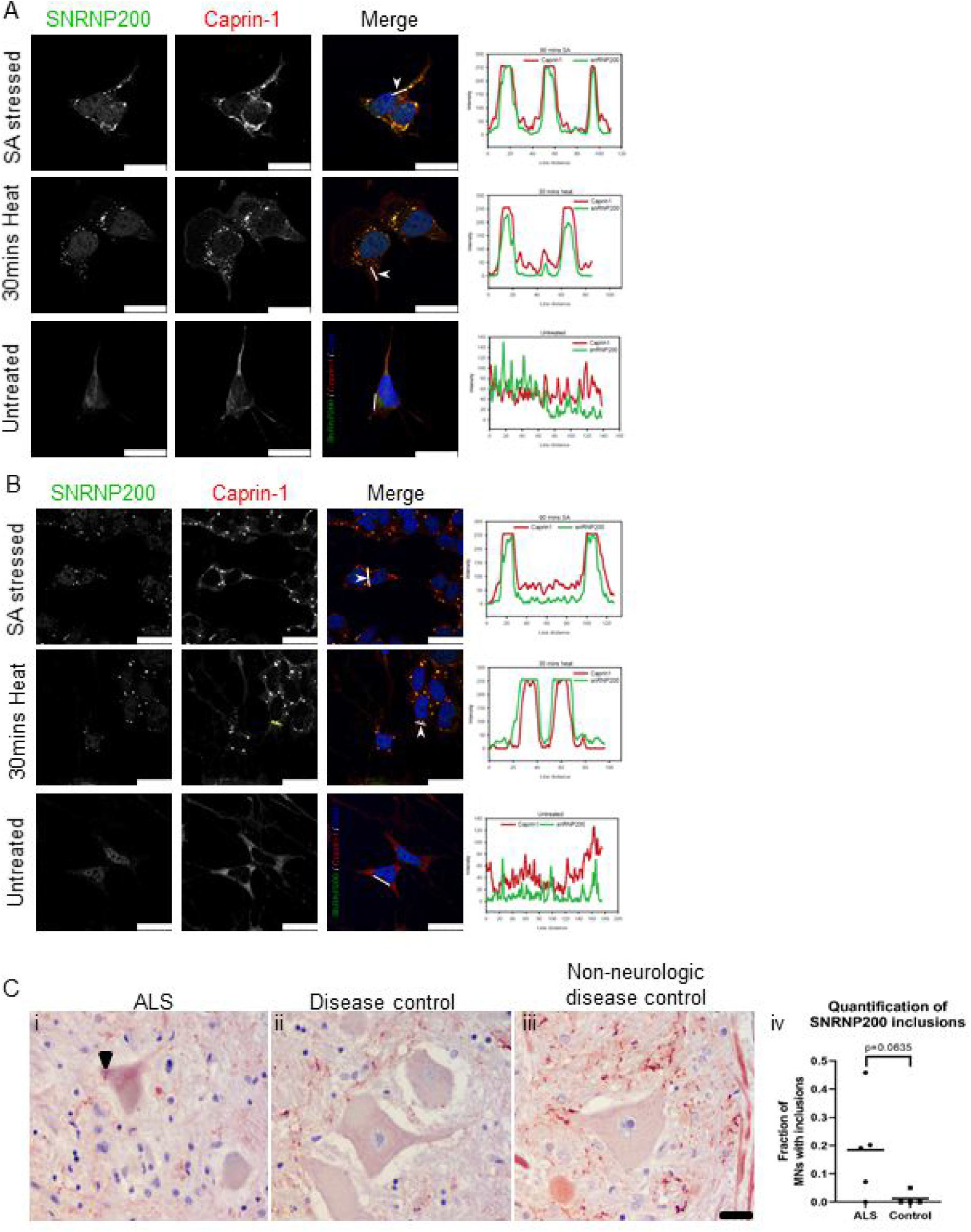
SNRNP200 co-localizes with SGs in neuronal-like cells and was exhibited in cytoplasmic inclusions in ALS post mortem spinal cord tissues. Representative images of SNRNP200 with Caprin-1 co-labelling in A) undifferentiated or B) differentiated SH-SY-5Y neuronal-like cells following SA (0.5mM, 30 mins) or heat stress (43°C, 30 mins). Nuclei are visualized with Hoechst. Line scan analysis indicates colocalization. Scale bar, 25 μm. C) Immunostaining for SNRNP200 on post mortem spinal cord tissues from i) ALS, ii) disease controls and iii) non-neurologic disease controls. Scale bar, 25 μm. Black arrowhead denotes SNRNP200 cytoplasmic inclusion. D) Quantification of SNRNP200 cytoplasmic inclusions in ALS and controls. Mann Whitney U test was used to assess statistical differences between the two groups.

### SNRNP200 is found in aggregates in post-mortem ALS neurons

Our *in vitro* studies demonstrated that SNRNP200 co-localized with Caprin-1 containing SGs in both HeLa as well as differentiated and undifferentiated SH-SY5Y cells in all stress conditions. Interestingly, SNRNP200 has been previously identified as an interactor of TDP-43 in HEK293 cells (39) and Caprin-1 has been reported as localized within cytoplasmic inclusions in sporadic ALS patient spinal motor neurons (11). Therefore, we speculated that SNRNP200 may also be re-distributed into cytoplasmic aggregates in motor neurons of ALS patients. To address this, we performed immunostaining on post-mortem lumbar spinal cord and motor cortex tissues from ALS cases, neurologic disease controls (DCs), and non-neurologic disease controls (NNDCs). SNRNP200 labeling was diffuse in most motor neurons of both DC and NNDC controls (Figure 6C). In contrast, SNRNP200 labeling was found as cytoplasmic inclusions within some motor neurons of both the lumbar spinal (Figure 6C, black arrowhead) and motor cortex (Figure S6, black arrowhead) of ALS cases. Quantification revealed a trend for the fraction of spinal cord motor neurons displaying SNRNP200 inclusions to be higher in ALS cases relative to controls, but did not reach statistical significance (Figure 6D, 18% in ALS vs. 1% in controls, p = 0.0635).

## Discussion

In this study, we investigated the Caprin-1 interactome to provide new insight into the proteome of SGs. Prior SG proteome studies were performed primarily using G3BP1 as the bait protein. These studies usually employ either overexpression (14) or the integration of tagged sequences (18, 19), each of which may perturb SG protein stoichiometry or structure. While these studies have provided tremendous insight, characterization of SGs at physiological levels is lacking. Our study addresses this gap through the IP of endogenous Caprin-1, providing insight into the native proteome of SGs. In addition to the stressed interactome, we also examined the unstressed interactome of Caprin-1. Our pathway analyses demonstrated that the interactors in both the stressed and unstressed conditions annotated to similar pathways. There was a considerable overlap (166 proteins; Table S3) between the unstressed and stressed interactors suggesting pre-stress interactions exist between Caprin-1 and ribosomal, RNA binding, and spliceosome proteins. Similar pre-stress interactions have also been observed with the G3BP1 interactome (18, 19). These pre-stress complexes may serve as seeds to facilitate rapid nucleation of SGs under stressed conditions, as previously proposed (18, 19). Interestingly, Caprin-1 interacts with G3BP1 in both stressed and unstressed conditions, therefore, it is reasonable to posit similarities in their interactomes. However, investigation of the overlap between our stress dependent Caprin-1 interactor list and three published G3BP1 proteomics studies indicate minimal overlap (Figure S3; Table S7). These discrepancies may be due to the different cell lines and stress regimes used in the studies. Moreover, overexpression or proximity labeling approaches capture the most abundant or nearby interactors, respectively, which may also account for the minimal overlap found between our study and previously published G3BP1 interactomes. The interactions within RNP granules are dynamic and not all proteins interact within the same time scale during SG assembly and disassembly (19). Interactions captured during acute stress (30 mins) as used in the BioID labeling approaches (18) may be altered during the recovery phase, as performed in our study. In summary, probing the SG interactome at various time points will be required to capture the dynamism of SG-related PPIs. Moreover, the use of multiple bait proteins will be necessary to catalogue the complete SG proteome, as one study has recently reported (20).

From our IP-MS results, we selected 12 candidate RBPs to further validate as SG proteins, and four proteins (SNRNP200, ANKHD1, Talin-1, GEMIN5) co-localized with SA induced SGs in HeLa cells (Figures 4 and 5). Our IP-MS methodology used whole cell lysates, thus these data suggest that the other eight candidate RBPs may interact with Caprin-1 in diffuse cytoplasmic complexes. Future studies may modify our IP-MS protocol to fractionate the cell lysate similar to the protocol used to isolate SG cores (14). Only SNRNP200 co-localized with SGs in neuronal-like cells (Figure 6), highlighting that the SG proteome may differ between cell types or in vivo. Although a prior study has demonstrated GEMIN5 to be associated with EIF4G labelled SGs in both SA and thermal stress conditions in HeLa cells (40), we did not observe GEMIN5 co-localization with Caprin-1 following thermal stress. SNRNP200 was the only candidate that exhibited co-localization to SGs irrespective of the stressor and cell type used. To the best of our knowledge, there is no prior evidence of SNRNP200 localization to SGs.

ALS and FTLD are pathologically characterized by phosphorylated TDP-43 (pTDP-43) containing cytoplasmic inclusions in motor neurons and glia (41). Previous studies have suggested that TDP-43 cytoplasmic aggregates co-localize with SG markers in ALS and FTLD *postmortem* tissues (42). However, the protein composition of cytoplasmic inclusions in patient derived tissues remains unclear. Our study identified SNRNP200 as a novel SG protein in our *in vitro* data, and also a component of some cytoplasmic inclusions within the motor neurons of ALS patient spinal cord and motor cortex. Whether these inclusions are part of the pTDP-43 or p62 inclusions remain unclear and thus future studies are required to determine co-localization with SNRNP200. Recent evidence reported downregulation of *SNRNP200* gene expression in ALS spinal cord and motor cortex relative to controls (43). Collectively, these results are suggestive of a functional depletion of SNRNP200 protein in affected motor neurons in ALS patients, possibly further indicating splicing defects in ALS.

In addition to some SNRNP200 labelling of cytoplasmic inclusions, we also observed a punctate staining pattern within the parenchyma and along the periphery of motor neurons in both the ALS and control cases used in this study. Prior work in rats has demonstrated RNG105 (ortholog of human Caprin-1) granule structures that accumulate near post-synaptic terminals in the dendrites of hippocampal neurons (36). Additionally, another study profiled the postsynaptic density protein 95 (PSD95) interactome in the anterior cingulate cortex and determined that SNRNP200 was among the list of identified interactors upregulated in bipolar patients relative to controls (44). We hypothesize that the SNRNP200 puncta we observed may be localized to pre/post synaptic terminals but further studies are required to address this hypothesis.

Taken together, we characterized the Caprin-1 protein interactome in both unstressed and stressed conditions. Our results highlighted that Caprin-1, a component not regulated by TDP-43 or G3BP1 (22), interacts with components of the ribosome, RNA transport proteins, spliceosome components in unstressed conditions. Additionally, these associations persist and coalesce into SGs when subjected to various stressors. One Caprin-1 interactor, SNRNP200, is a novel SG component *in vitro* in neuronal and non-neuronal cells, and also exhibited cytoplasmic aggregation within the motor neurons of post-mortem ALS patient spinal cord and motor cortex. While the cause/consequence of these inclusions remains to be elucidated, these results highlight Caprin-1 SGs and its binding partners as potential contributors to altered RNA metabolism in neurodegenerative disease.

## Acknowledgements

We thank Patrick Dion and Jay Ross for important discussions and the CRCHUM Cellular Imaging platform. We gratefully acknowledge funding from the ALS Canada/Brain Canada Arthur J. Hudson Translational Team grant (CVV, RB) and the Target ALS Human *Postmortem* Tissue Core for access to human tissue samples. We thank Ray and Amy Thurston for their generous funding contribution (PP). Research reported in this publication included work performed in the mass spectrometry core supported by the National Cancer Institute of the National Institutes of Health under grant number P30CA033572. The content is solely the responsibility of the authors and does not necessarily represent the official views of the National Institutes of Health.

## Supplementary Figure Legends

**Figure S1: Gene ontology analysis using DAVID on the unstressed interactors of Caprin-1.** A) The top 15 biological processes and B) top 15 Molecular functions terms. Benjamini-Hochberg correction was applied and all terms shown exhibited p <0.05.

**Figure S2: Known protein-protein interactions for SA-stress dependent Caprin-1 interactors.** StringDB analysis demonstrating protein-protein interactions between the stressed dependent interactors of Caprin-1 with distinct clusters annotated to the RNA binding proteins (pink) and splicing factors (blue). All other interactors are labeled in gray.

**Figure S3: Comparison of stress-dependent Caprin-1 interactome with previously published SG interactomes.** A) Comparison with BioID-based Caprin-1 interactome in HEK293 cells following acute SA stress for 30 mins. B) Comparison with BioID-based G3BP1 interactome of SGs in HEK293 cells following acute SA stress for 30 mins. C) Comparison with SG interactome in U2OS cells probed by a GFP-tagged G3BP1 following acute SA stress for 30 mins. D) Apex-G3BP1 labelling based SG interactome in HEK293T cells following SA stress for 1 hr.

**Figure S4: Shortlisted Caprin-1 interactors which did not co-localize with SGs in HeLa cells subjected to oxidative stress.** HeLa cells were either stressed with 0.5mM SA (left panel) or unstressed (right panel) and co-stained with Caprin-1 and the respective candidate proteins. Nuclei were visualized with Dapi. Scale bar, 25μm.

**Figure S5: ANKHD1, Talin-1, and GEMIN5 did not co-localize with Caprin-1 labelled SGs in neuronal-like cells.** SH-SY5Y cells were stressed with either or 0.5mM SA (left) or 30 mins heat at 43°C (middle) or left untreated (right) and co-labelled for Caprin-1 and A) ANKHD1, B) Talin-1, and C) GEMIN5. Nuclei were visualized with Hoechst. Scale bar, 25 μm. D) Differentiated SH-SY5Y cells were labelled with Tuj1 and Hoechst. Scale bar, 50 μm.

**Figure S6: SNRNP200 detected as cytoplasmic inclusions in ALS post mortem motor cortex tissues.** A) ALS, B) disease controls, and C) non-neurologic disease controls. Scale bar, 25 μm. Black arrowhead denotes SNRNP200 cytoplasmic inclusion.

## Supplementary Tables

**Table S1:** Antibodies used for immunostaining experiments.

**Table S2:** Table of all proteins identified from each IP experiment and their overlap from Figure 1D.

**Table S3:** Proteins list corresponding to the Venn diagram of high confidence interactors as predicted by the SAINTexpress analysis from Figure 2A.

**Table S4:** List of unstressed Caprin-1 interactors that were annotated to KEGG pathways from Figure 2B.

**Table S5:** List of stress dependent Caprin-1 interactors that were annotated to KEGG pathways from Figures 3A-B.

**Table S6:** Proteins list corresponding to the Venn diagrams from Figure S3.

**Table S7:** Proteins list corresponding to the Venn diagram comparing RNA binding proteins and significant high confidence Caprin-1 interactors from Figure 3C.

