## Supplementary figures and images for "Defining the Caprin-1 interactome in unstressed and stressed conditions"

### Supplemental Figures

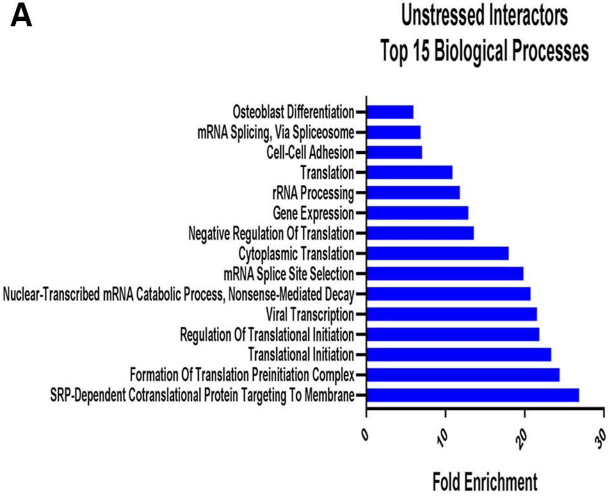

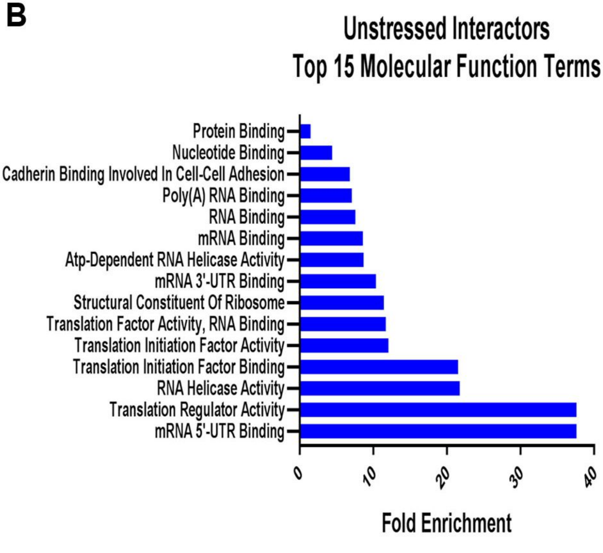

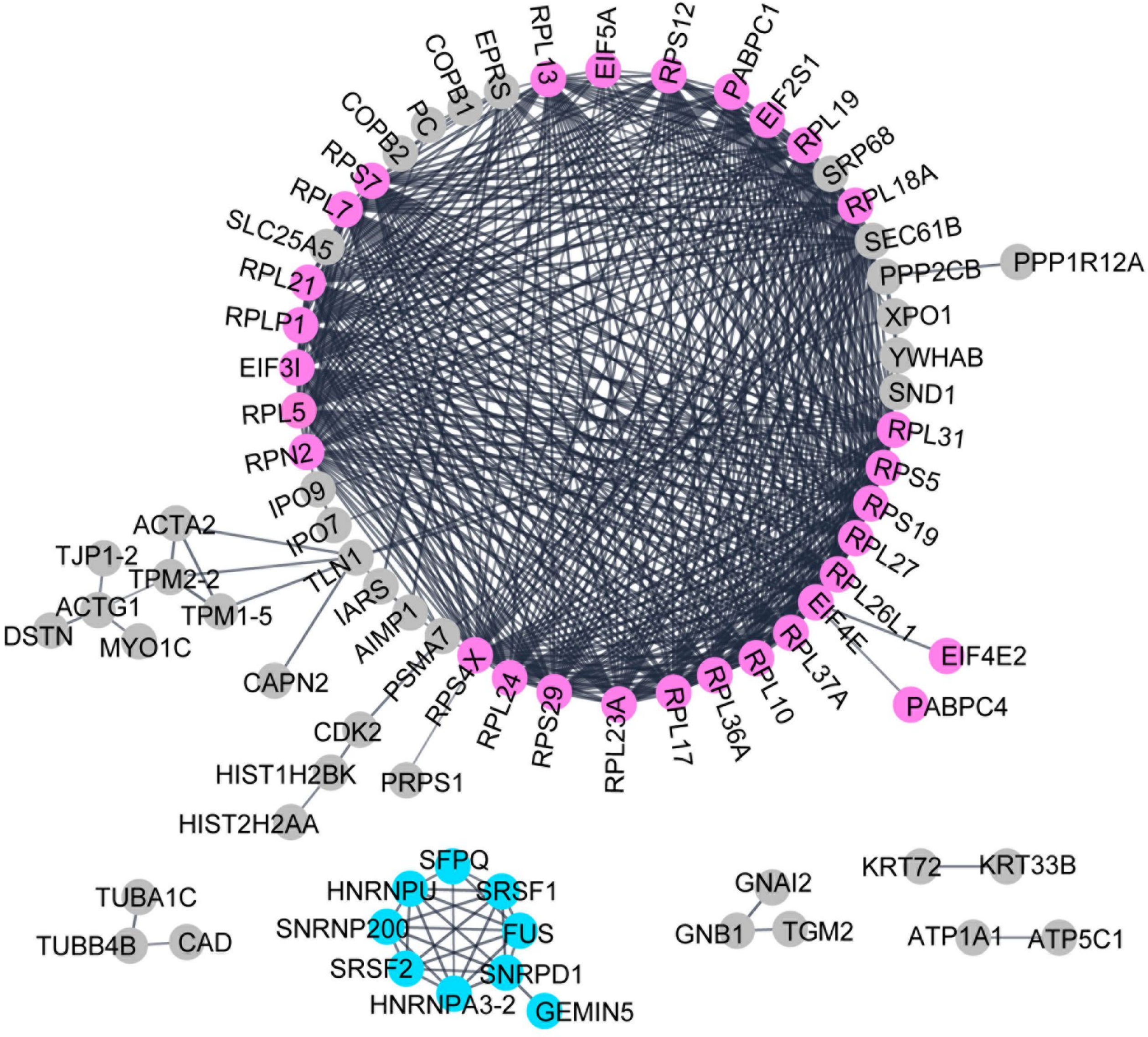

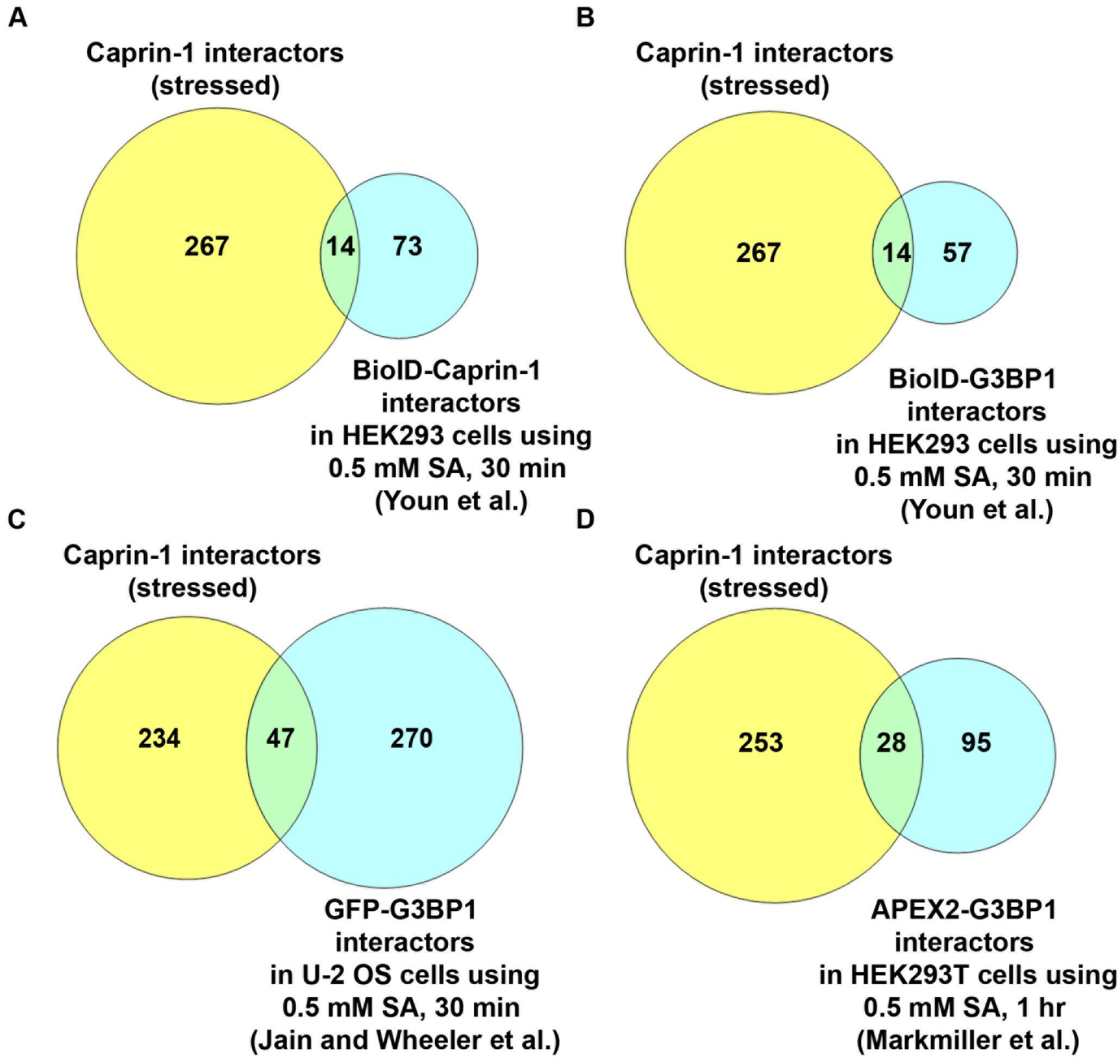

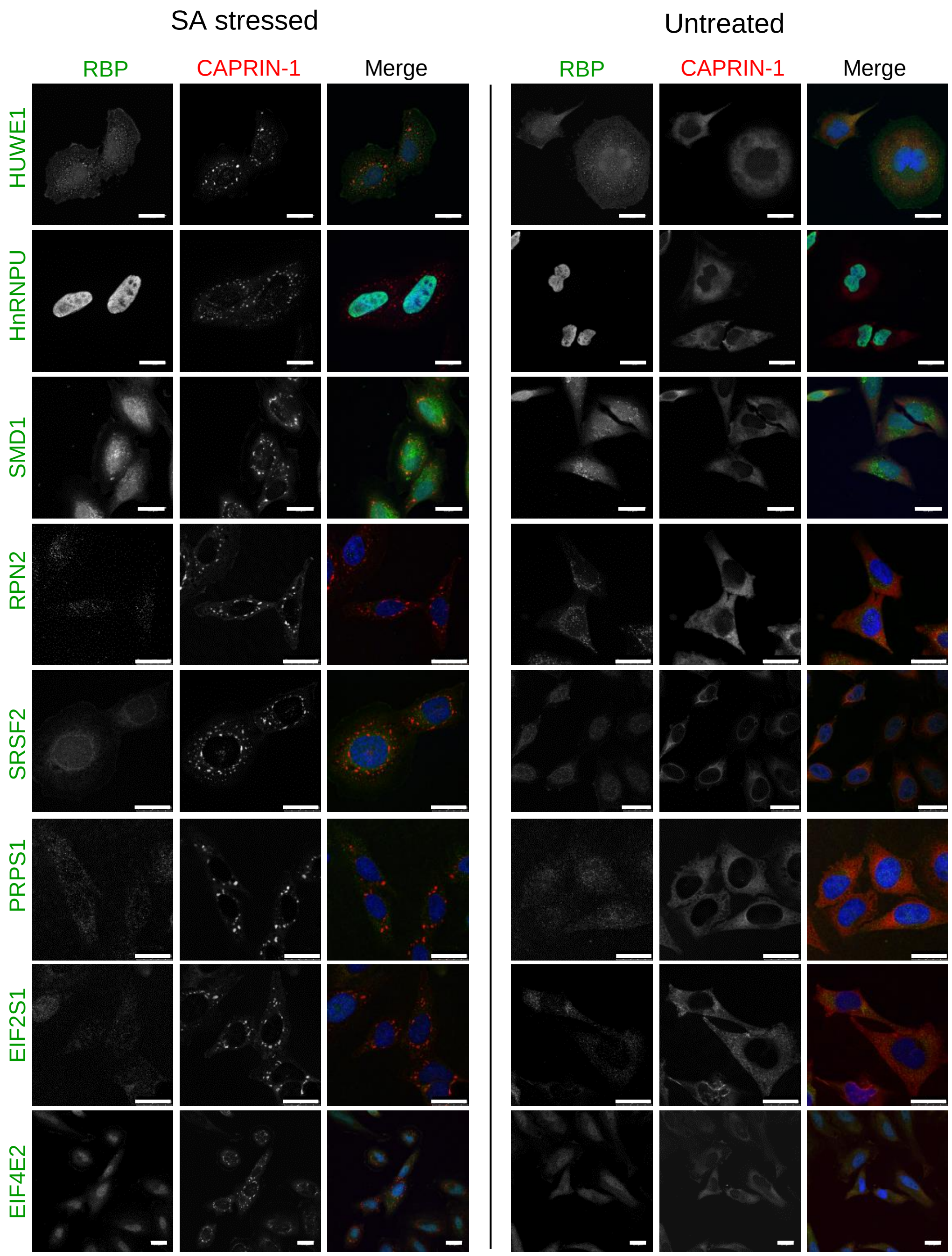

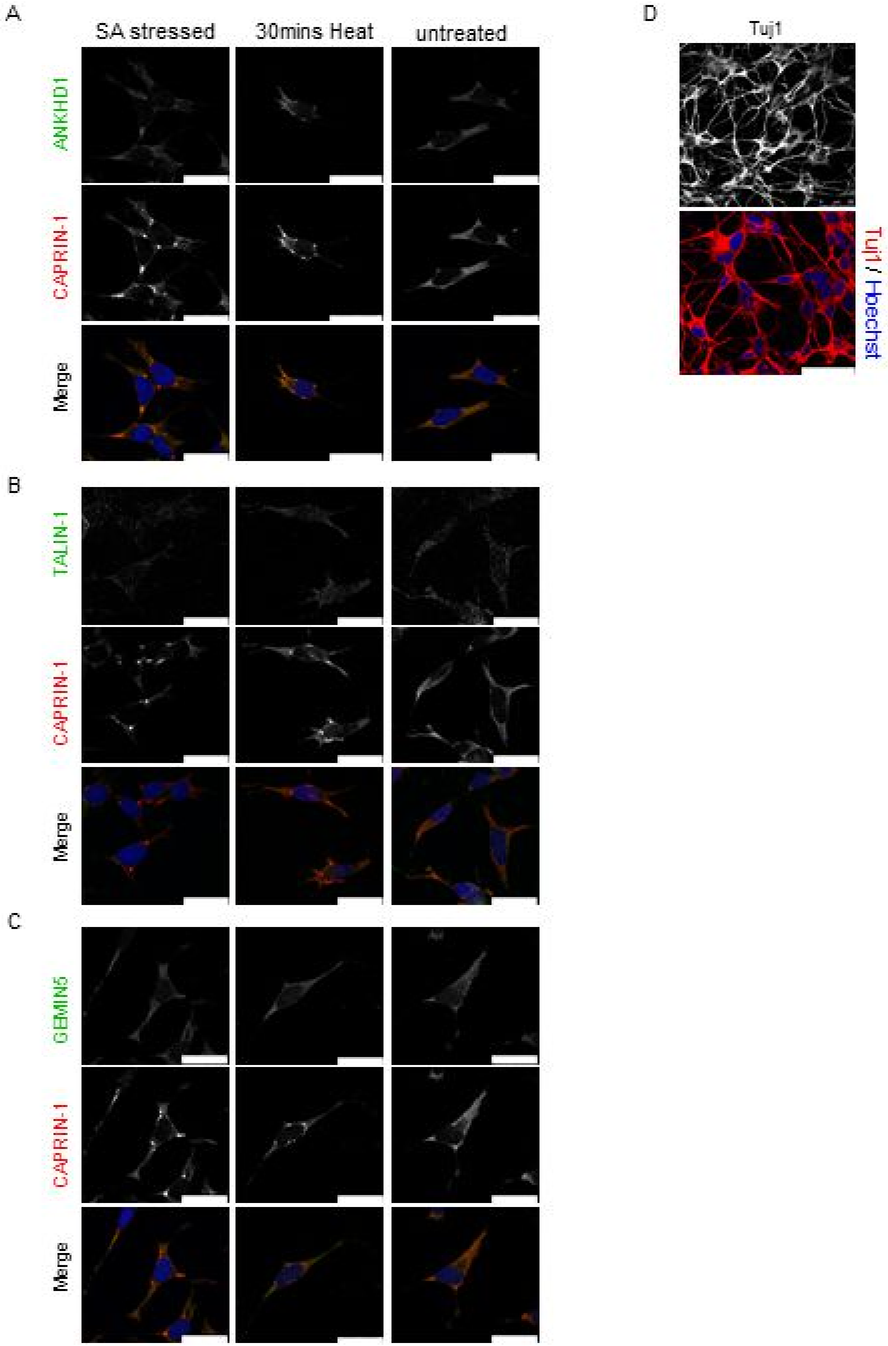

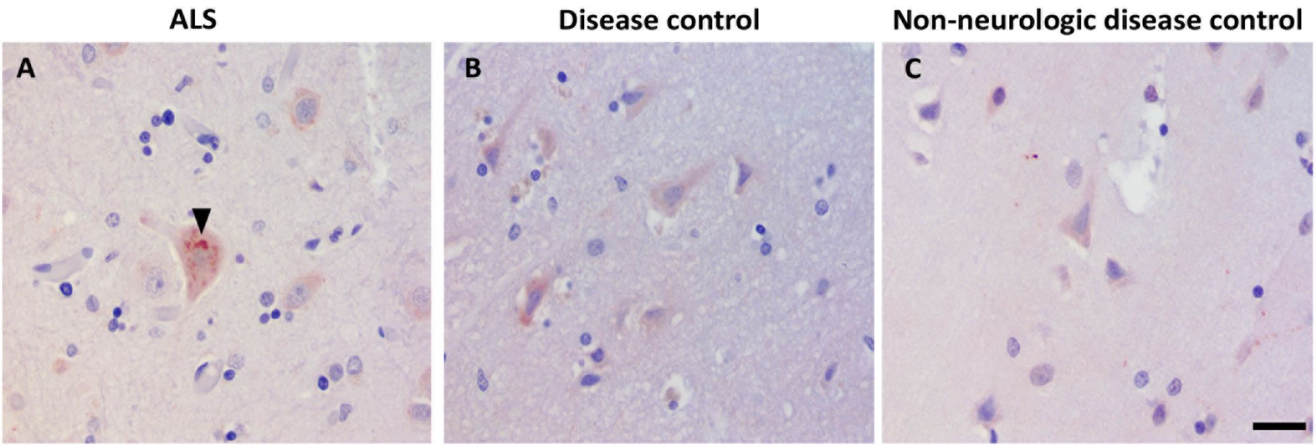

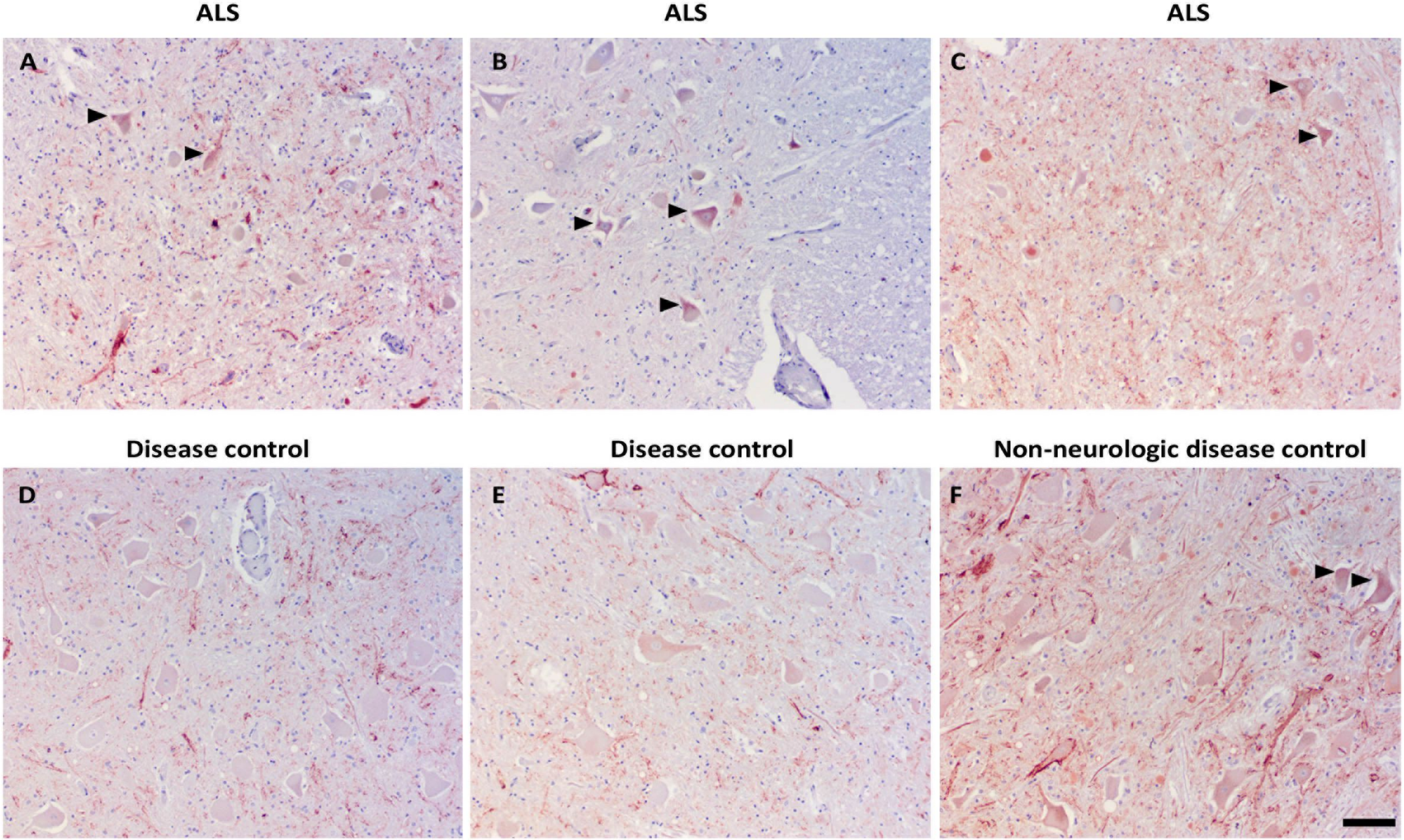
